# Loss of KRAS^G12D^ feedback regulation involving splicing factor SRSF1 accelerates pancreatic cancer

**DOI:** 10.1101/2021.10.13.464210

**Authors:** Ledong Wan, Kuan-Ting Lin, Mohammad Alinoor Rahman, Zhikai Wang, Mads A. Jensen, Youngkyu Park, David A. Tuveson, Adrian R. Krainer

## Abstract

The gene encoding KRAS GTPase is recurrently mutated in pancreatic ductal adenocarcinoma (PDAC), triggering the formation of precursor lesions, i.e., acinar-to-ductal metaplasia (ADM) and pancreatic intraepithelial neoplasia (PanIN). However, the majority of pancreatic cells from KC (*LSL-Kras^G12D/+^; Pdx-1-Cre*) mice expressing the *Kras^G12D^* mutation remain morphologically normal for a long time, suggesting the existence of compensatory feedback mechanisms that buffer aberrant *Kras^G12D^* signaling, and that additional steps are required for disrupting cell homeostasis and promoting transformation. Here we report a feedback mechanism in which the ubiquitously expressed splicing factor SRSF1—which is associated with cell transformation in multiple cell types—is downregulated in the majority of morphologically normal pancreas cells with the *Kras^G12D^* mutation. Conversely, increasing SRSF1 expression disrupts cell homeostasis by activating MAPK signaling, in part by regulating alternative splicing and mRNA stability of interleukin 1 receptor type 1 (*Il1r1*). This disruption in homeostasis in turn accelerates *Kras^G12D^*-mediated PDAC initiation and progression. Our results demonstrate the involvement of SRSF1 in the pancreatic-cell homeostatic response against the *Kras^G12D^* mutation, dysregulation of which facilitates PDAC initiation.

**One-Sentence Summary:** Splicing factor SRSF1 is involved in KRAS^G12D^ feedback regulation and pancreatic-cancer tumorigenesis.

## Main Text

Extensive efforts, including retrospective reconstructions of clonal evolution from deep DNA sequencing of tumors, have provided insights into tumor evolution. However, fundamental gaps still exist in our understanding of early-stage tumor initiation. In particular, numerous somatic mutations accumulate in healthy cells during the course of aging, yet they require a long time to evolve into carcinoma and do so with relatively low frequency (*1–3*). The current view of tumor evolution is that genetic and epigenetic alterations are required for tumorigenesis, in addition to driver-oncogene mutations. Moreover, negative feedback mechanisms counteract oncogenic activation, contributing to oncogene-induced senescence and termination of oncogenic signals (*4–6*). However, how cells can remain physiologically normal, despite having oncogene mutations, remains unclear. A better understanding of the homeostatic responses of cells against driver-oncogene mutations should facilitate early cancer diagnosis and the development of new therapies.

Pancreatic ductal adenocarcinoma (PDAC) has an extremely poor prognosis, attributable to the presence of distant metastases at the time of diagnosis, which limits traditional cancer treatments like surgery or chemotherapy (*7*). Understanding the process of PDAC initiation is especially critical for PDAC prevention and effective treatment. Somatic activating mutation in *KRAS* is present in >90% of PDAC, and is dominated by substitution from glycine to aspartate (G12D) at the 12th residue, within the G domain (*8, 9*). Studies involving the genetically engineered KC mouse model (*LSL-**K**ras^G12D/+^*; *Pdx1-**C**re*) support the notion that the *Kras^G12D^* mutation triggers acinar-to-ductal metaplasia (ADM) and pancreatic intraepithelial neoplasia (PanIN), and results in a protracted onset of PDAC (*9–11*). However, the majority of pancreatic cells from two-month-old KC mice remain morphologically normal (Fig. 1A) (*9*), even though they harbor an activated *Kras^G12D^* allele, with a single LoxP site detected by PCR after recombination of the “lox-stop-lox (LSL)” cassette (fig. S1, A and B). To examine the excision-recombination of the LSL cassette *in situ*, we designed two sets of RNA fluorescent *in situ* hybridization (FISH) probes complementary to sense and antisense strands of the LSL cassette, respectively (fig. S1A). Notably, the majority of pancreas acinar cells had a recombined LSL cassette, with loss of the FISH signal corresponding to an active *Kras^G12D^* allele (fig. S1, C and D). These observations suggest the existence of a compensatory feedback mechanism against mutant *Kras^G12D^*, such that additional alterations are presumably required to overcome this homeostatic response.

**Fig. 1.**
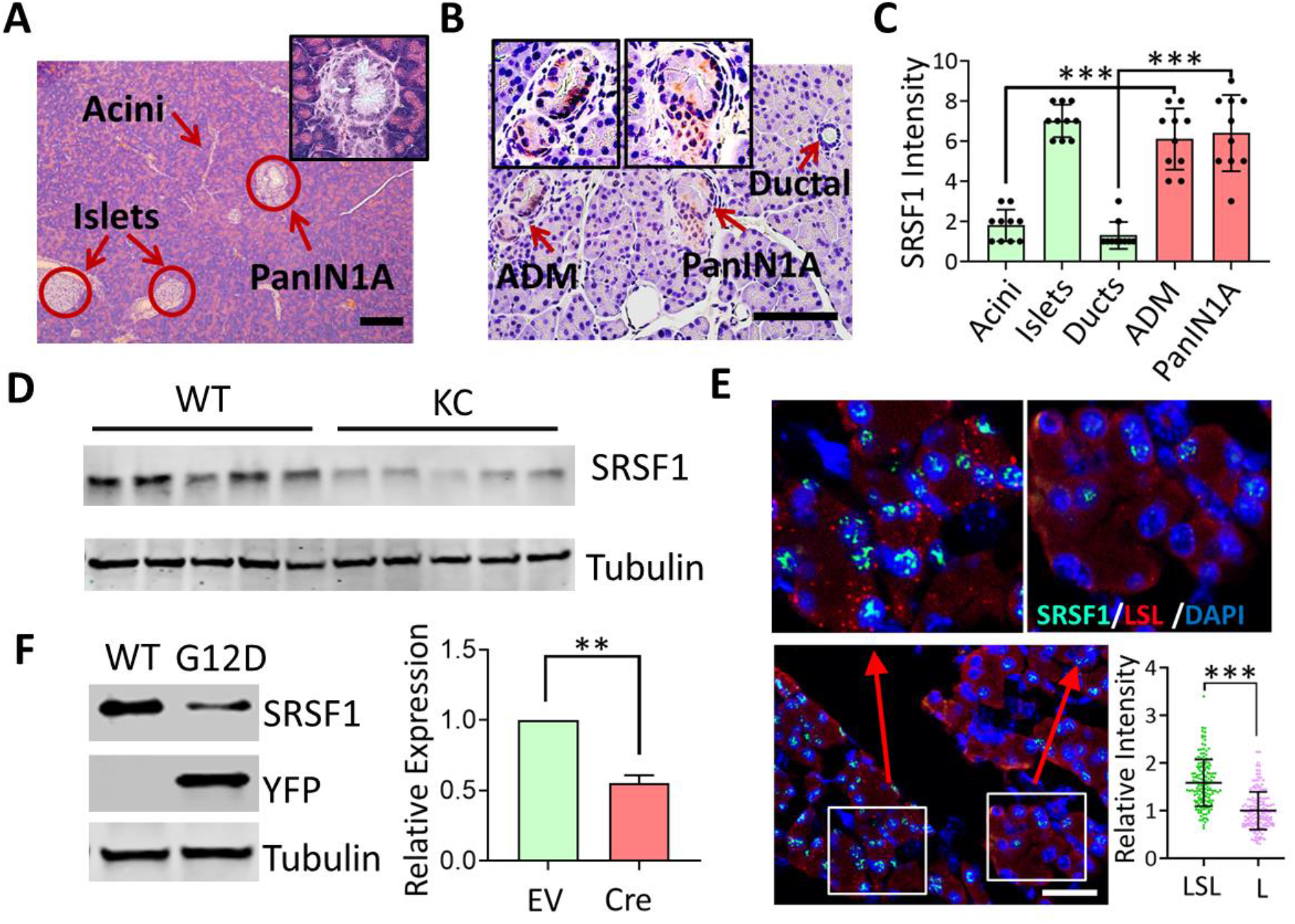
Reduced expression of SRSF1 in morphologically normal *Kras^G12D^* pancreatic cells, and increased expression in PanIN lesions. (**A**) Hematoxylin and eosin (H&E) staining of pancreata from two-month-old KC mice. Inset shows higher magnification of PanIN1A lesion. Scale bars, 100 μm. (**B**) SRSF1 IHC staining of pancreata from two-month-old KC mice. Insets show higher magnification of ADM and PanIN1A lesions. Scale bars, 100 μm. (**C**) IHC intensity scores of SRSF1 in different pancreas cells, ADM, and PanIN lesions (n = 10 per group). One-way ANOVA with Tukey’s multiple comparison test. \*\*\**P* < 0.001. (**D**) SRSF1 protein levels in pancreatic protein lysates from WT (wild-type) and KC mice (n = 5 per group). (**E**) Representative images of multiplexed FISH/IF staining of STOP-cassette and SRSF1 protein in pancreata isolated from KC mice. Bottom right, quantification of SRSF1 intensity in LSL-positive or -negative pancreas cells. (**F**) Immunoblot (left) and quantification of SRSF1 protein (right) of *LSLKras^G12D/+^*; *R26-LSl-YFP* ductal organoids infected with Adeno-Empty (WT) or Adeno-Cre (G12D) (n = 3 biological replicates). (E, F) Unpaired, two-tailed t test. \*\**P* < 0.01, \*\*\**P* < 0.001. Error bars represent mean ± SD.

Dysregulation or mutation of oncogenic splicing factors contributes to tumor initiation and progression by altering pre-mRNA alternative splicing events that impinge on multiple signaling pathways (*12, 13*). Slight overexpression of the splicing factor SRSF1 (formerly SF2/ASF) proto-oncogene promotes transformation and tumorigenesis, e.g., in fibroblast and mammary-epithelial cell contexts (*14, 15*). We report that SRSF1 mRNA and protein are upregulated in both human and mouse PDAC, compared to normal tissue (fig. S2, A to E). Elevated expression of SRSF1 is associated with poor prognosis in PDAC patients (fig. S2, G and H). Conversely, SRSF1 knockdown in the human PDAC SUIT-2 cell line inhibited proliferation and metastasis, suggesting that elevated SRSF1 contributes to PDAC maintenance (fig. S3).

Consistent with its role in cell transformation, we detected elevated SRSF1 protein in both ADM and PanINs in KC mice, as well as in the context of cerulein-induced pancreatitis and ADM (*11, 16*) (Fig. 1, B and C, fig. S2F and fig. S4). These results suggest that elevated SRSF1 is involved in PDAC initiation, including the onset of ADM.

Given the pro-transformation role of SRSF1, we next characterized its expression in the context of the homeostatic response to *Kras^G12D^*. We harvested the pancreas from early-disease-stage KC mice and *Pdx1-Cre* control littermates. In contrast to its elevated expression in ADM, PanIN and PDAC, SRSF1 protein decreased in the pancreas of two-month-old KC mice (Fig. 1D). Due to the mosaic expression of Cre recombinase in KC mice, a proportion of acinar cells retain an intact LSL cassette and a dormant *Kras^G12D^* allele (fig. S1D) (*9*). Therefore, we combined FISH and immunofluorescence staining to examine the expression pattern of SRSF1 in KC mice *in situ*. Remarkably, SRSF1 expression was reduced in the population of morphologically normal acinar cells with an active *Kras^G12D^* allele, compared with those harboring a dormant *Kras^G12D^* allele (Fig. 1E).

To confirm that SRSF1 downregulation participates in the response of normal cells to *Kras^G12D^*, we generated normal ductal organoids from *LSL-Kras^G12D/+^*; *R26-LSL-YFP* mice. Both LSL cassettes repressing *Kras^G12D^* and YFP were then excised by adenoviral-Cre infection (fig. S5, A to C). Consistent with our *in vivo* data, we observed a significant decrease in SRSF1 expression upon acute activation of the *Kras^G12D^* allele (Fig. 1F). The observed decrease of SRSF1 in the morphologically normal cells with active *Kras^G12D^* suggests that SRSF1 downregulation is part of the compensatory feedback to the *Kras^G12D^* mutation.

To further examine the association of SRSF1 with cell transformation, we generated a mouse model dubbed SC (*tetO-**S**rsf1*; *LSL-rtTA*; *Pdx-1-**C**re*) with doxycycline-inducible expression of SRSF1 in the pancreas (fig. S6A). Dox-treated SC mice exhibited histologic signs of pancreatitis, including interstitial edema, lymphocyte infiltration, collagen deposition, and accumulation of metaplastic ductal lesions (Fig. 2A). Flow cytometry revealed an influx of immune cells into the pancreas, especially macrophages (Fig. 2B and fig. S7A), which we confirmed by immunofluorescence staining (fig. S7B). Moreover, we detected elevated serum levels of the pancreatic enzymes amylase and lipase upon short-term Dox treatment, and these enzymes returned to their normal levels after 30 days of SRSF1 induction, consistent with the transition from acute to chronic pancreatitis (Fig. 2, C and D) (*16*). Additionally, SRSF1-induced pancreatitis was reversible after a recovery period following Dox withdrawal (fig. S8), suggesting that SRSF1 contributes to the duration of inflammation. Together with the observed increase in SRSF1 in the cerulein-induced ADM lesions (fig. S4B), these data demonstrate that elevated SRSF1 induced pancreatitis and ADM.

**Fig. 2.**
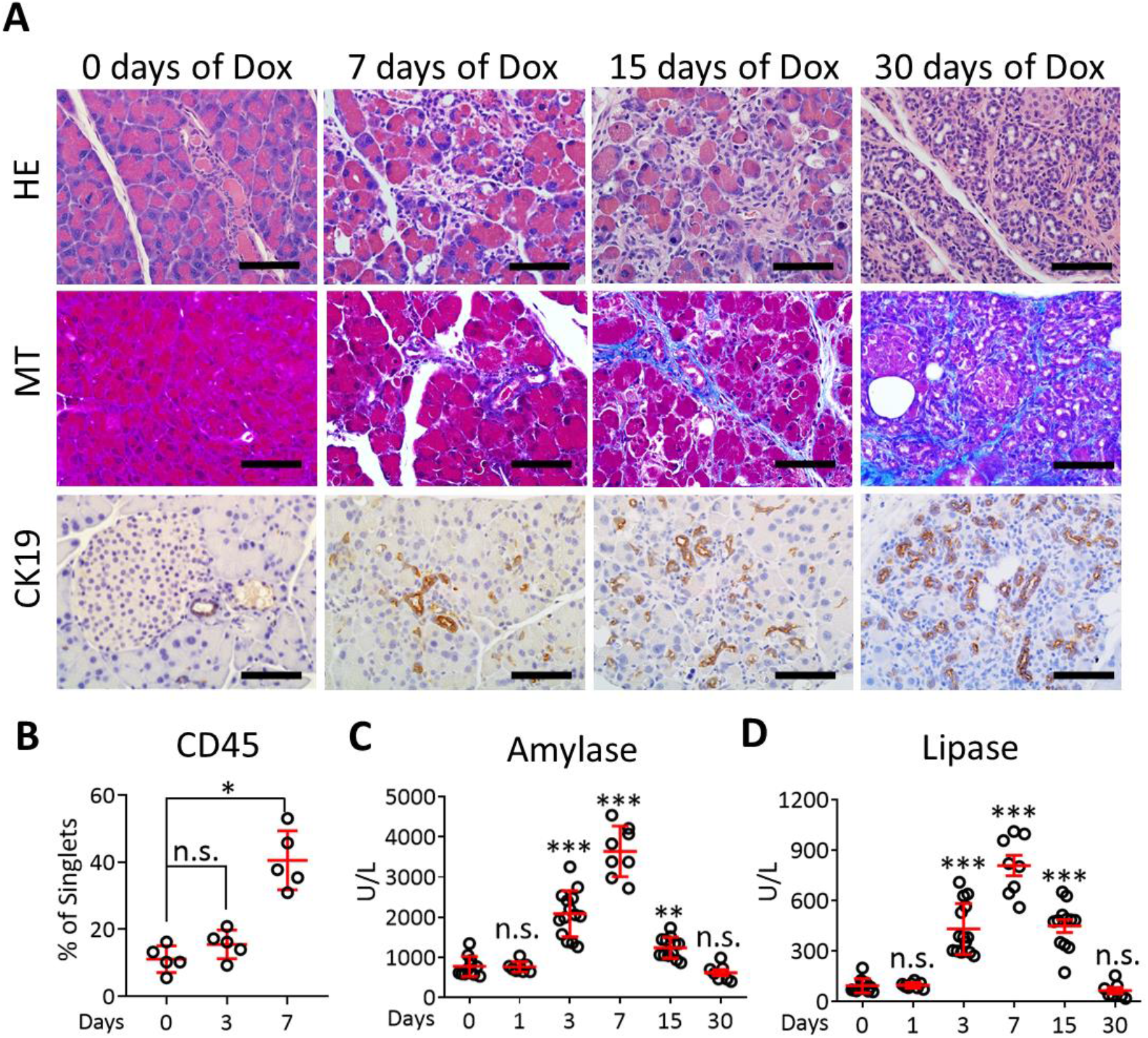
SRSF1 promotes pancreatitis and ADM. (**A**) Histological evaluation of SC mice by H&E staining, Masson’s trichrome staining (MT) (blue indicates collagen deposition), and cytokeratin CK19 expression by IHC to identify metaplastic ductal lesions, after treatment with Dox. Scale bars, 50 μm. (**B**) Immune-cell infiltration evaluated by flow cytometry in SC mice treated with Dox. (**C**, **D**) Circulating levels (U/L) of the pancreatic enzymes amylase (C) and lipase (D) in SC mice after treatment with Dox (days). Each data point represents a measurement from an individual mouse (B - D) Kruskal-Wallis test followed by pairwise comparisons using Wilcoxon rank-sum test. Family-wise error rate was adjusted using Bonferroni-Holm method. \**P* <0.05, \*\**P* < 0.01, \*\*\**P* <0.001. Error bars represent mean ± SD.

To investigate whether elevated SRSF1 can disrupt homeostasis in cells with *Kras^G12D^* and promote tumorigenesis, we generated the KSC strain by intercrossing SC and *LSL-Kras^G12D/+^* mice (Fig. 3A and fig. S6B). Compared to the pancreatic parenchyma of two-month-old KC mice, which was mostly morphologically normal, with rare low-grade PanINs, the vast majority of the pancreatic cells in KSC mice were transformed and exhibited neoplasia (Fig. 3B and fig. S9). In addition, KSC mice exhibited extensive, high-grade PanIN lesions, characterized by loss of cell polarity, significant nuclear atypia, and budding of cell clusters into the ductal lumen (Fig. 3, A and C). Next, we further examined the effect of SRSF1 in PDAC development by introducing a conditional *LSL-Trp53^R172H^* allele, generating the KPSC strain (fig. S6C). Even though KPC mice (*LSL-**K**ras^G12D/+^*; *LSL-Tr**p**53^R172H/+^*; *Pdx-1-**C**re*) exhibit advanced PDAC with markedly shortened median survival, compared to KC mice *(10)*, we detected only rare low-grade PanINs in one-month-old KPC mice. In contrast, KPSC mice rapidly succumbed to PDAC tumors, with even shorter median survival (68 days) than KPC mice (139 days) (Fig. 3, D and E, and fig. S10). We conclude that SRSF1 contributes to PDAC initiation and progression in both KSC and KPSC contexts.

**Fig. 3.**
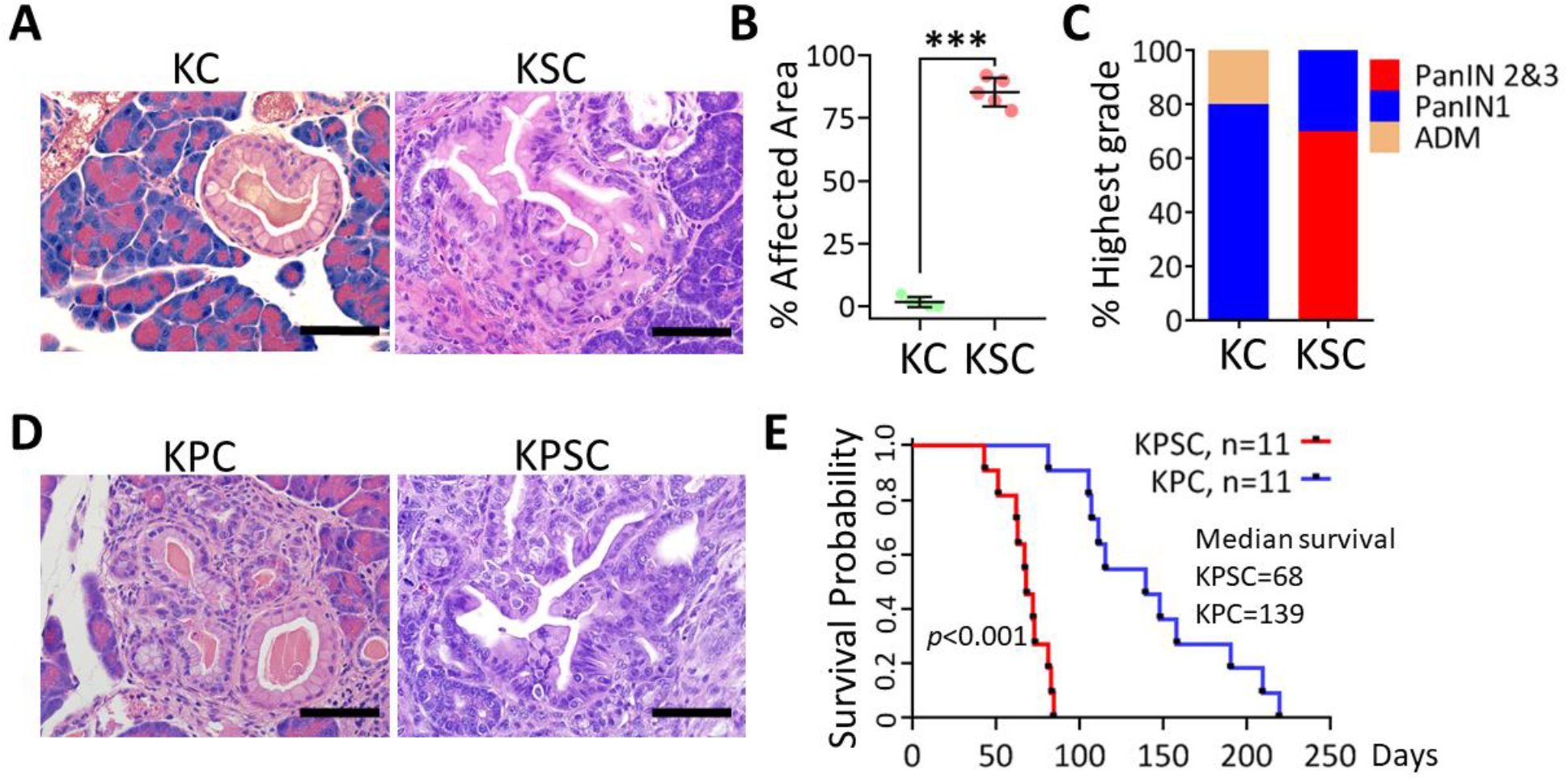
SRSF1 accelerates *Kras^G12D^*-mediated tumorigenesis. (**A**) H&E staining of pancreata from two-month-old KC and KSC mice. Scale bars, 100 μm. (**B**) Quantification of the percentage of pancreatic area exhibiting histological signs of transformation and neoplasia from two-month-old KC and KSC mice (n = 5 per group). Unpaired two-tailed t test, \*\*\**P* <0.001. Error bars represent mean ± SD. (**C**) Classification of highest-grade lesions present in two-month-old KC and KSC mice (n = 5 per group). (**D**) H&E staining of pancreata from one-month-old KPC and KPSC mice. Scale bars, 100 μm. (**E**) Kaplan-Meier survival analysis of KPC and KPSC mice (n = 11 per group). The *P* value was determined by a log-rank Mantel-Cox test.

To address the underlying mechanisms of how decreased SRSF1 contributes to the homeostatic response to *Kras^G12D^*, and conversely, how elevated SRSF1 accelerates *Kras^G12D^*-mediated disease progression, we generated organoids from SC and KSC mice, and these organoids exhibited comparable induction of SRSF1 after Dox treatment (fig. S11A). Pathway-enrichment analysis showed negative enrichment of the MAPK signaling pathway in the presence of the *Kras^G12D^* mutation (Fig. 4A). This result is consistent with compensatory feedback in response to RAS activation, which dampens RAS signaling output (*4, 5*). In accord with previous reports (*17, 18*), only the subset of cells with hyper-activated MAPK signaling exhibited pancreatic metaplasia and neoplasia, even within individual ducts from two-month-old KC mice (fig. S11B). These results point to the pivotal role of MAPK signaling in cell homeostasis, and the need for this pathway to be activated in the context of cell transformation. Remarkably, induction of SRSF1 consistently strengthened MAPK signaling in both wild-type and *Kras^G12D^* mutant contexts (Fig. 4, A and B, and fig. S11C), demonstrating that SRSF1 is involved in the cellular homeostatic response to *Kras^G12D^*, and that increasing SRSF1 accelerates *Kras^G12D^*-mediated tumorigenesis by modulating MAPK signaling output.

**Fig. 4.**
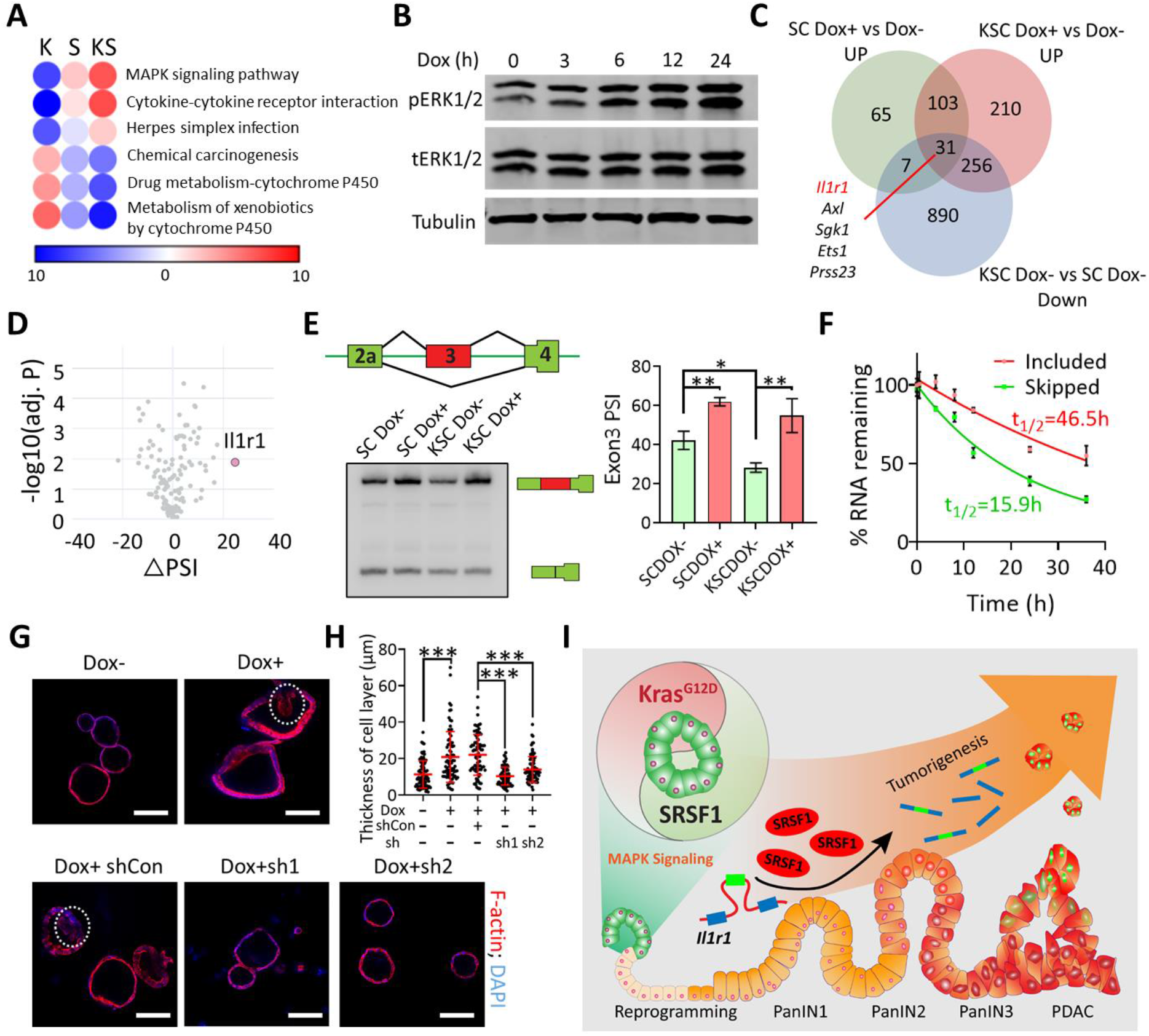
Loss of regulation of SRSF1-IL1R1-MAPK signaling promotes *Kras^G12D^*-mediated PDAC initiation. (**A**) Pathway-enrichment analysis of genes with increased (red) or decreased (blue) expression, in KSC organoids compared to SC organoids without Dox treatment (K), Dox-treated SC organoids compared to untreated SC organoids (S), and Dox-treated KSC organoids compared to untreated KSC organoids (KS). Color bar represents −log10 (*P* value). (**B**) SC organoids were evaluated by immunoblotting for activation of the MAPK signaling pathway after treatment with Dox. (**C**) Venn diagram of genes up-regulated in SC and KSC organoids after treatment with Dox; and genes down-regulated in KSC organoids compared to SC organoids without Dox treatment. (**D**) Volcano plots of aberrant splicing events in the MAPK signaling pathway from SC organoids after treatment with Dox. (**E**) Schematic and radioactive RT–PCR results of SRSF1-regulated splicing event in *Il1r1* pre-mRNA. The percent spliced in (PSI) was quantified for each condition (n = 3 biological replicates). (**F**) mRNA-decay assay of the *Il1r1* isoforms in SC organoids at the indicated times after actinomycin D treatment. mRNAs were quantified by RT-qPCR, normalized to *Gapdh* levels, and expressed as a percentage of the levels at time 0 h (n = 3 biological replicates). (**G**) Representative IF staining for F-actin (red) with DAPI counterstain of the indicated KSC organoids treated with Dox, and infected with control (shCon) or *Il1r1* shRNAs. Dashed circles indicate multiple cell layers. Scale bars, 100 μm. (**H**) Quantification of the thickness of cell layers of the indicated KSC organoids treated with Dox, and transduced with control (shCon) or *Il1r1* shRNAs. (**I**) Model of the SRSF1 decrease contributing to the homeostatic response to *Kras^G12D^* mutation in normal cells, followed by the SRSF1 increase, which disrupts MAPK-signaling homeostasis, accelerating *Kras^G12D^*-mediated tumorigenesis. \**P* <0.05, \*\**P* < 0.01, \*\*\**P* <0.001 by one-way ANOVA with Tukey’s multiple comparison test. Error bars represent mean ± SD.

Importantly, we observed that multiple MAPK-signaling-pathway genes were both upregulated upon SRSF1 overexpression and downregulated in response to *Kras^G12D^* (Fig. 4C and fig. S12A). Aberrant pre-mRNA splicing of MAPK signaling components has been implicated in tumorigenesis (*15, 19–21*). Interestingly, among the genes with opposite expression changes in response to elevated SRSF1 and *Kras^G12D^*, we identified interleukin1 receptor type 1 (*Il1r1*) as the top SRSF1-regulated splicing target associated with the MAPK signaling pathway (Fig. 4D and fig. S12B). SRSF1 consistently promoted inclusion of exon 3 in the 5’ untranslated region of *Il1r1* mRNA, in SC and KSC organoids (Fig. 4E).

*Il1r1* encodes a cytokine receptor for IL1α, IL1β, and IL1RN, activation of which leads to the production of inflammatory mediators and the regulation of biological responses that include tissue vascularity, adipogenesis, lipid metabolism, and inflammation (*22–25*). Suppression of IL1R1 with a proprietary antagonist, Anakinra, attenuates the level of pERK and suppresses PDAC tumor growth (*26*). We found elevated IL1R1 within pancreatitis lesions (fig. S13A), as well as in the PanIN lesions in KC mice (fig. S13C), consistent with the reported association of IL1R1 with PDAC progression (*27*). Moreover, we observed increased IL1R1 in response to SRSF1 overexpression in SC and KSC mice (fig. S13, B and C). To address the relationship between the SRSF1 effects on *Il1r1* expression and on pre-mRNA splicing, we measured the decay rate of the two mRNA isoforms, and found that inclusion of alternative exon 3 increases *Il1r1* mRNA stability (Fig. 4F). Furthermore, an *Il1r1* minigene assay showed enhanced inclusion of exon 3 upon overexpression of SRSF1 (fig. S14, A to C). To test the dependence of exon 3 inclusion on direct SRSF1 binding to *Il1r1* pre-mRNA, we mutated two SRSF1 motifs in exon 3 (fig. S14A). As expected, mutating the motifs individually or simultaneously suppressed SRSF1-induced exon 3 inclusion (fig. S14, B and C). We conclude that SRSF1 binds to exon 3 in the *Il1r1* pre-mRNA and promotes IL1R1 expression via alternative-splicing-regulated mRNA stability.

Next, we assessed the contribution of IL1R1 to SRSF1-mediated cell transformation by knocking down IL1R1 in the organoid model. SRSF1 overexpression resulted in a change in KSC organoid morphology, characterized by the development of multiple-cell-layer features (*28*), and this abnormal phenotype was rescued by IL1R1 knockdown (Fig. 4, G and H, and fig. S15).

These results demonstrate that SRSF1 promotes pancreatitis and ADM, and coordinates the *Kras^G12D^*-driven cellular homeostatic response and tumorigenesis by fine-tuning MAPK signaling (Fig. 4I). Specifically, *Il1r1* expression decreased upon the homeostatic response to *Kras^G12D^* mutation, and increased upon SRSF1-regulated alternative splicing contributing to mRNA stabilization, consistent with the pivotal contribution of IL1R1 to MAPK signaling in PDAC (*26*). Of note, the recombinant IL1R1 antagonist Anakinra is being evaluated in PDAC clinical trials, in combination with standard chemotherapy (NCT02021422, NCT02550327, NCT04926467). Besides MAPK signaling, pathway-enrichment analysis also identified other pathways—including Cytokine-cytokine receptor interaction—with opposite regulation in response to *Kras^G12D^* and SRSF1 (Fig. 4, A and C, and fig. S12A), implying cross-interactions with the microenvironment in PDAC initiation. Our own and others’ previous studies indicate that SRSF1 expression is regulated via complex transcriptional (via MYC), post-transcriptional, and translational mechanisms (*29–33*). In particular, we found a strong positive correlation between *MYC* and *SRSF1* expression at the RNA level in PDAC tumors (fig. S16A), and also found that SRSF1 translation is regulated by translation-initiation factors, e.g. eIF4A and eIF4E (fig. S16B). Therefore, it will be of interest to determine the precise mechanisms underlying the SRSF1 expression changes involved in the homeostatic response to *Kras^G12D^*, and how they become dysregulated in tumorigenesis. In addition to its well-studied role in alternative-splicing regulation, SRSF1 has been implicated in other biological processes, e.g., translation, miRNA processing, protein sumoylation, and the nucleolar stress response (*34*), which may also be involved in cell homeostasis and PDAC development. Elevated SRSF1 is thought to contribute to multiple types of cancer (*34*), some of which are associated with various *KRAS* mutations (*35*). Comprehensive investigation of the relationship between SRSF1 and other *KRAS* mutations may enhance our understanding of the context-dependence of the cellular homeostatic response. Finally, the current study identified SRSF1’s involvement in the early cellular response to *Kras^G12D^*, and revealed that dysregulation of this response accelerates PDAC tumorigenesis; further understanding of the underlying mechanisms may identify early PDAC diagnostic markers and suggest new therapeutic strategies.

## Acknowledgments

We acknowledge the assistance of the Cold Spring Harbor Laboratory (CSHL) Animal and Genetic Engineering, Animal and Tissue Imaging, Microscopy, Flow Cytometry, Antibody, Next-Generation Sequencing, and Histology Shared Resources. We thank R. Jaenisch (Whitehead Institute) for KH2 ES cells; J. E. Wilkinson (University of Michigan) for pathology consulting; T. Ha (CSHL) for biostatistics consulting; and R. Karni (Hebrew University of Jerusalem), H. H. Zhang, and J. L. Tang (Zhejiang University) for helpful discussions. The TROMA III (CK19) antibody was obtained from the Developmental Studies Hybridoma Bank, created by the NICHD of the NIH and maintained at The University of Iowa. Access to the PanCuRx dataset was authorized by the Ontario Institute for Cancer Research, supported by the Government of Ontario.

## Author contributions

Conceptualization: LW, ARK. Methodology: LW, YP, DAT, ARK

Investigation: LW, KTL, MAR, ZW, MAJ, YP Writing: LW, ARK

Resources: DAT, ARK Supervision: ARK

## Data and materials availability

RNA-seq data is available at the Sequence Read Archive of NCBI under the BioProject ID: PRJNA747810.

## Supplementary Materials

Materials and Methods

Figures S1-S16

Tables S1

References (*36–48*)

